# Optimizing high-yield production of SARS-CoV-2 soluble spike trimers for serology assays

**DOI:** 10.1101/2020.05.27.120204

**Authors:** Dominic Esposito, Jennifer Mehalko, Matthew Drew, Kelly Snead, Vanessa Wall, Troy Taylor, Peter Frank, John-Paul Denson, Min Hong, Gulcin Gulten, Kaitlyn Sadtler, Simon Messing, William Gillette

## Abstract

The SARS-CoV-2 spike trimer is the primary antigen for several serology assays critical to determining the extent of SARS-CoV-2 exposure in the population. Until stable cell lines are developed to increase the titer of this secreted protein in mammalian cell culture, the low yield of spike protein produced from transient transfection of HEK293 cells will be a limiting factor for these assays. To improve the yield of spike protein and support the high demand for antigens in serology assays, we investigated several recombinant protein expression variables by altering the incubation temperature, harvest time, chromatography strategy, and final protein manipulation. Through this investigation, we developed a simplified and robust purification strategy that consistently yields 5 mg of protein per liter of expression culture for two commonly used forms of the SARS-CoV-2 spike protein. We show that these proteins form well-behaved stable trimers and are consistently functional in serology assays across multiple protein production lots.

**Highlights:**

- Improved yields of SARS-CoV-2 spike protein through modification of expression and purification parameters
- Yields of greater than 5 mg/l obtained for VRC spike under optimal conditions
- Spike protein quality was validated by QC methods to ensure utility in serology assays

## 1. Introduction

The need for high quality protein reagents is an important aspect of the response to the COVID-19 pandemic [1]. Screening for the presence and extent of an immune response will be a critical tool in controlling the spread of the infection until the development of an effective vaccine. Additionally, such monitoring will provide essential data to policy makers as they develop guidelines to limit virus spread in the population. With the ubiquitous presence of related coronaviruses in the population (e.g. SARS-CoV, MERS-CoV, common cold coronaviruses OC43 and HKU1), the tests must be highly specific for SARS CoV-2. Several serology assays, both published and in progress, are employing the S protein (hereafter referred to as spike) in ELISA-based assays. The specificity inherent in the spike protein [1,2] makes it an obvious target for therapeutic interventions and for use in serology studies to assess the prevalence of immune responses to a specific coronavirus. Two spike antigens are currently widely used: the receptor binding domain (RBD), which interacts with the extracellular ACE2 receptor, and a much larger soluble spike ectodomain modified for stability to mimic the prefusion native spike trimer conformation [2]. While robust production of spike RBD domain was recently reported [3], high quality soluble spike trimers remain a difficult antigen to both express and purify. Publication of one form of this protein cites a yield of 0.5 mg/l [2], with another publication suggesting yields as high as 5 mg/l [3]. However, many unpublished reports indicate consistent yields only in the 1-2 mg/l range, providing a significant challenge for large-scale serology assay development. In addition, some of these publications provide limited information on purification details or protein quality assessment, which make interpretation of the yields challenging.

To support an NIH-led serosurvey on the extent of the coronavirus immune response in ∼ 10,000 human samples, our lab was tasked to produce RBD and spike proteins for ELISA assay optimization and deployment [4]. We found our RBD production yield to be similar to published reports, however, spike production was problematic with our initial attempts producing inconsistent results and lower yields than those reported in the literature. Without a robust method in place in our lab for determining spike titer in culture supernatants, we were limited to the crude method of SDS-PAGE gel analysis for estimating expression levels. Due to the size of spike (138 kDa), heavy level of glycosylation, and low expression levels, SDS-PAGE gel analysis was inconclusive when comparing multiple lots of expression supernatants. Thus, the work presented here is intended to provide a robust method for those wishing to reliably produce SARS CoV-2 spike protein in quantities sufficient for serology assays, structural biology, or simply to better understand some of the production variables affecting the yield. It is expected that this protocol could be further improved and perhaps eventually replaced by a stable cell line production platform. Nevertheless, the approaches outlined here allowed us to improve the production yield of spike protein significantly by modifying cell culture temperature and harvest time, as well as improving the purification process. The final proteins produced were highly pure, formed appropriate trimeric structures, and were functional as antigens in ELISA assays. Taken together, these improvements allowed the production of sufficient spike antigen for more than 500 ELISA plates per liter of culture and generated enough protein from a single 4-liter expression to support a robust serosurvey being conducted by the NIH [4].

## 2. Materials and methods

### 2.1 DNA

DNA for the expression of VRC spike (SARS-CoV-2 S-2P-3C-His8-Strep2×2) and spike proteins for SARS-CoV, MERS-CoV, OC43-CoV, and HKU1-CoV were generously provided by Dr. Kizzmekia Corbett (Dale and Betty Bumpers Vaccine Research Center, NIAID). DNA for the expression of Mt. Sinai spike (SARS-CoV-2 S-2P-His6) was generously provided by Dr. Florian Krammer (Icahn School of Medicine, Mt. Sinai) through BEI Resources. The similarities and differences in these constructs are outlined in **Fig. 1**. Transfection-quality DNA was produced in-house using the Qiagen Plasmid Plus Maxi Kit per the manufacturer’s protocols or was generated at large-scale by Aldevron (Fargo, ND).

**Figure 1.**
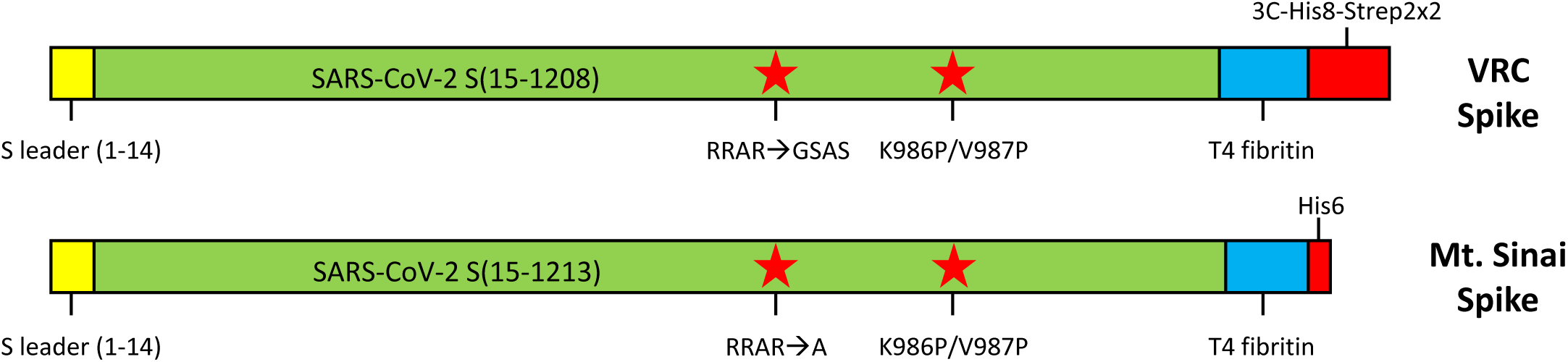
Comparison of VRC and Mt. Sinai spike protein expression constructs. Both constructs contain the native spike (S) protein signal sequence (amino acids 1-14) followed by the S ectodomain (green). Furin cleavage sites (RRAR) were mutated in both constructs as noted, and stabilizing proline mutations were introduced. Both constructs contain a phage T4 fibritin trimerization domain (blue) and C-terminal purification tags (red). The Mt. Sinai construct contains a non-cleavable His6 purification tag, while the VRC construct contains a combination His8-dual Strep2 tag preceded by a rhinovirus 3C protease cleavage sequence.

### 2.2 Mammalian cell culture

Manufacturer’s protocols were followed for the transfection and culturing of Expi293F cells (Thermo Fisher Scientific, Waltham MA). Briefly, 1.7 liters of cell culture at 2.9 x 10^6^ cells/ml were transfected with preformed Expifectamine:DNA complexes at 1 μg/ml of final culture volume. Expression cultures were incubated at 37°C and 8% CO_2_ in 5-liter Optimum Growth Flasks (Thomson Instrument Company, Oceanside, CA) shaking at 105 RPM on an Infors HT Multitron Standard shaker with a 2” orbit. Expression enhancers were added 18-20 hours post-transfection per manufacturer’s instructions, and incubation was continued at 37°C and 8% CO_2_ until harvest time of either 72, 96, or 120 hr post-transfection. For temperature shift experiments, the incubation temperature was lowered to 32°C immediately after enhancer addition.

### 2.3 Tangential flow filtration (TFF)

Harvested culture supernatants were clarified by centrifugation (4000 x *g*, 20 min, 4°C) followed by filtration (catalog# 12993, Pall Corporation, Port Washington, NY). When used in column chromatography, clarified supernatants were concentrated and buffer exchanged by TFF. Specifically, a MasterFlex peristaltic pump (Vernon Hills, IL) fed the clarified supernatant to a 30 kDa MWCO cassette (catalog# CDUF002LT, MilliporeSigma, Burlington, MA). The clarified supernatant was concentrated to 10% of the initial volume and then buffer exchanged with 5 volumes of 1x PBS, pH 7.4 (Buffer A, diluted from 10X PBS, catalog #70011069 Thermo Fisher Scientific, Waltham, MA). Following buffer exchange, the TFF cassette was rinsed with 200-250 ml Buffer A, to collect any protein remaining in the cassette. Clarified supernatants for use in batch purification were not buffer exchanged.

### 2.4 Protein purification

Chromatography was conducted at room temperature (∼22°C) using NGC medium-pressure chromatography systems from BioRad Laboratories Inc. (Hercules, CA) with exceptions noted below. The standard purification protocol employed immobilized metal affinity chromatography (IMAC) followed by desalting into final buffer. Specifically, TFF-treated culture supernatant was adjusted to 25 mM imidazole and applied to a 5 ml Ni Sepharose High Performance nickel-charged column (GE Healthcare, Chicago, IL) previously equilibrated in Buffer A + 25 mM imidazole. The flow rate for all steps of the IMAC was 5 ml/min. TFF-processed culture supernatant from up to 8 liters of cell culture was applied to a single 5 ml column. The column was washed in Buffer A + 25 mM imidazole for 4 column volumes (CV) with the final 3 CV collected separately as the column wash. The protein was eluted from the column by first reversing the orientation of the column, applying a 6 CV linear gradient of 25 mM – 175 mM imidazole in Buffer A, a 6 CV step elution of Buffer A + 325 mM imidazole, and a 3 CV step elution of Buffer A + 500 mM imidazole. 2.5 ml fractions were collected for all elution steps. Elution fractions were analyzed by SDS-PAGE/Coomassie-staining and appropriate fractions were pooled. Typical pool volume for a purification from four liters of culture supernatant was ∼30 ml.

The sample was desalted into Buffer A using a HiPrep 53 ml 26/10 desalting column (GE Healthcare, Chicago, IL) with 14 ml injections at 9 ml/min for all steps. The final protein sample was created by combining the bulk elutions from multiple runs of the desalting column. The protein concentration was determined by measuring the A_280_ using a Nanodrop One spectrophotometer (Thermo Fisher Scientific, MA, USA). Final protein was dispensed as 0.5 ml and 0.05 ml aliquots, snap frozen in liquid nitrogen, and stored at -80°C. To assess the oligomeric state of the protein, a single 0.5 ml aliquot of the final protein was thawed and analyzed by analytical size-exclusion chromatography using a 10/300 Superdex200 analytical column (GE Healthcare, Chicago, IL), with a flow rate of 0.5 ml/min.

For the batch purification from filtered (see above) culture supernatants, 10 ml of Ni-charged MagBeads (GeneScript, Piscataway, NJ), previously equilibrated in Buffer A, were placed on the bottom of 2 x 5-liter Thompson flasks (5 ml per flask) and filtered culture medium was added the flasks (1 to 2 liters/flask depending on the expression culture volume). Flasks were shaken at 105 rpm at 27°C for 3 hr. After incubation, the supernatant was decanted into a four liter glass beaker, a rare-earth magnet was used to capture the beads to the bottom of the beaker, and the medium was removed and saved as “flow through”. 300 ml of Buffer A was added to the flask to suspend the beads, which were transferred to 500 ml Corning centrifuge bottles and shaken at ∼22°C (room temperature for all subsequent steps) for 5 min. Beads were collected, the wash removed, and a second 300 ml wash step was performed. The beads were washed an additional two times with 40 ml of Buffer A in a 50 ml Corning tube while shaking for 5 min for each wash (total of 4 wash steps). Proteins were eluted from the washed beads by addition of 20 ml Buffer A + 25 mM imidazole for elution 1, followed by 5 elutions with 20 ml of Buffer A + 500 mM imidazole. 20 ml of appropriate buffer was used for each elution and the beads shaken in 50 ml Corning tubes for 10 min before collecting. Samples of each elution fraction were analyzed by SDS-PAGE and Coomassie-staining and appropriate fractions were pooled. The pool was treated as above for buffer exchange, final gel analysis, and storage.

### 2.5 Transmission electron microscopy (TEM)

Transmission electron microscopy of the purified VRC and Mt. Sinai spike proteins was carried out by dilution of thawed final samples to 0.02 mg/ml in 20 mM Tris-HCl, pH 8.0, 200 mM NaCl followed by loading onto glow-discharged carbon support film grids (CF200-CU, Electron Microscopy Sciences). Grids were washed twice in buffer (20 mM Tris-HCl, pH 8.0, 200 mM NaCl) and stained with 0.75% w/v uranyl formate (pH 4.5) three times using filter paper to blot away the stain prior to immediate application of additional stain. A final staining step was carried out for 30 sec with the stain then removed by wicking with filter paper, and the grids were dried under an incandescent lamp. Stained grids were imaged on a Hitachi 7650 electron microscope at 40,000x magnification.

### 2.6 Enzyme-linked immunosorbent assay (ELISA)

In order to assess batch-to-batch reproducibility of spike antigens, purified VRC spike proteins from five different purifications were used as antigens in an ELISA with positive control CoV-2 serum at multiple dilutions. The ELISA protocol was carried out as reported [4] using 100 ng per well of the various spike proteins and dilutions of 1:500, 1:10,000, and 1:100,000.

## 3. Results and Discussion

To produce SARS-CoV-2 antigens for the development of serology assays, we initially followed standard procedures for secreted protein production: transfection using the manufacturer’s protocols, expression at 37°C, harvest at three days post-transfection, tangential flow filtration of the culture supernatant, immobilized metal ion chromatography with linear gradient elution, and size exclusion chromatography. However, this process resulted in very low yields (0.9 and 0.3 mg/l, respectively, for Mt. Sinai and VRC spike as seen in **Table 1**). These yield levels were not high enough to support large serology studies without significant scaleup and associated high costs. The urgent need and the narrow time frame to provide protein support for NIH serology studies limited our ability to perform extensive troubleshooting and optimization. Rather, we assessed several parameters with internally controlled experiments (e.g. splitting an IMAC pool in two and passing the two aliquots over different SEC resins). This was essential as safety protocols for laboratory personnel during the pandemic limited the number of staff in the lab. Using this approach, we were able to obtain information on the effect of multiple parameters including the temperature of cell culture, harvest time, and mode of target capture.

**Table 1.** Effect of expression and purification parameters on spike yield. Unless noted, the post-harvest production process followed the procedure of TFF > IMAC > desalting column. Numbers represent yields from independent experiments. nd - not determined.

| Condition | VRC (mg/l) | Mt. Sinai (mg/l) |
| --- | --- | --- |
| 37°C/72 hr IMAC/SEC | 0.3 | 0.9 |
| 37°C/96 hr IMAC/desalt | 2.0, 2.0, 1.4 | 1.7, 2.6 |
| 32°C/96 hr IMAC/desalt | 4.8, 5.2, 6.1 | 4.6, 5.3 |
| 32°C/120 hr IMAC/desalt | 6.4, 5.0 | 4.4 |
| 32°C/96 hr MagBeads/desalt | 4.1, 5.5 | nd |

### 3.1 Effect of purification strategy

Inspection of the SDS-PAGE/Coomassie staining analysis of the initial IMAC (not shown), suggested that we could improve the separation of spike from contaminant by adjusting the elution parameters. The relatively high affinity of the His-tagged spike proteins for the column enabled the use of a 6 CV linear elution gradient from 25 mM to 175 mM imidazole which eluted the majority of contaminants before spike was eluted in a step elution of 325 mM imidazole (**Fig. 2A**). Analysis of intermediate purification steps by the low-resolution method of SDS-PAGE/Coomassie staining suggested significant protein loss was occurring during SEC. This correlated with published methods of spike purification [3] and suggested that high concentrations of spike might lead to protein loss by precipitation. We found it difficult to quantitate the amount of target loss during these steps, likely due to heavy glycosylation, which is reported to interfere with Bradford analysis of protein concentration [5,6]. By using a desalting column for buffer exchange, rather than SEC or diafiltration, our modified protocol is designed to minimize protein manipulations that might lead to protein loss especially during post-IMAC steps. Thus, we compared purification yields from parallel experiments with either SEC or a desalting column as the final purification step (**Table 1**). The desalting column protocol led to a consistently higher yield of ∼2 mg/l, indicating that this method improved protein recovery. In addition, very little difference in protein quality was observed in these samples, particularly in the case of the VRC spike protein, again arguing against the need for the SEC step. As can be seen in **Fig. 2B**, the VRC spike protein was consistently of higher purity than the Mt. Sinai spike protein. However, ELISA data comparing these two proteins suggests that these minor impurities have no significant effect on the antigenicity in the assay [4]. Nevertheless, for downstream processes which require higher levels of purity, Mt. Sinai spike may require additional purification steps.

**Figure 2.**
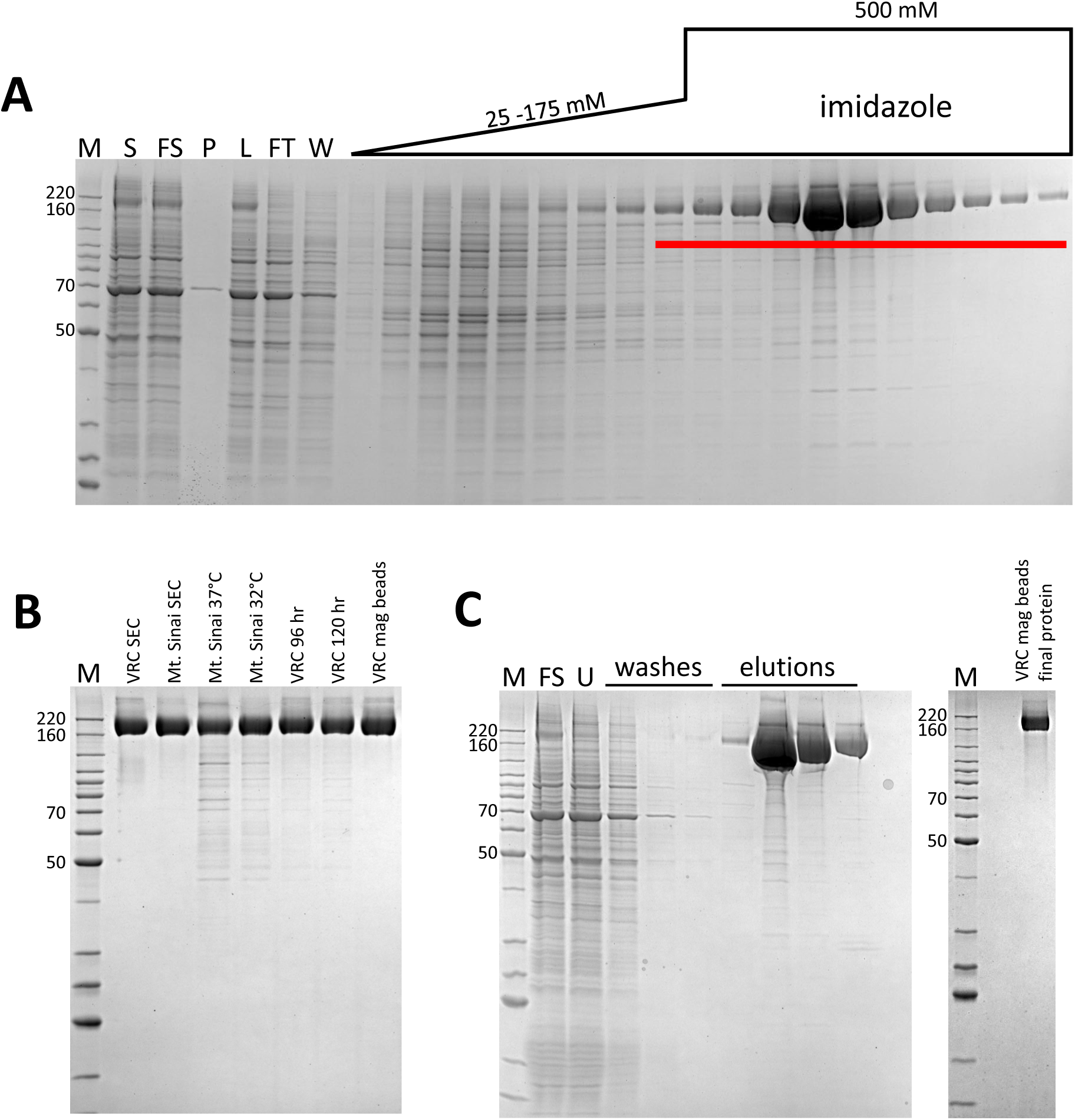
Representative IMAC chromatography fraction analyses and purified proteins. M – protein standards, molecular weights of select standards noted in kDa. **A.** Coomassie-stained SDS-PAGE analysis of standard IMAC protocol (VRC spike), S – culture supernatant, FS – filtered culture supernatant, P – TFF permeate, L – TFF retentate/column load, FT – column flow through, W – column wash. Fractions pooled are underlined. **B.** Coomassie-stained SDS-PAGE analysis of representative purified proteins. **C.** Representative Coomassie-stained SDS-PAGE analysis of Ni magnetic bead chromatography and final protein (VRC spike). FS – filtered culture supernatant, U – unbound protein.

### 3.2 Effect of expression temperature and time on spike production yield

Reducing expression temperature during recombinant protein expression, in both *E. coli* and the baculovirus/insect expression system, can significantly increase yields of soluble protein [7]. In transient CHO systems, temperature reductions are commonly used to enhance protein production [8]. However, similar modifications in HEK293-based systems are less frequently reported. Previous work in our lab and others suggests one limitation to protein secretion in transiently transfected mammalian cell culture might be the secretion process itself. While multiple factors appear to be involved, a recent paper suggests that limiting components in the secretory pathway appear to be the major bottleneck [9,10]. We did observe a significant amount of spike protein in the Expi293 cell pellets (data not shown), suggesting that some protein was being held up in the endoplasmic reticulum, or was otherwise failing to completely mature through the secretory pathway. Low cell viabilities (<70%) at harvest were another sign of potential toxicity caused by secretory failures. Thus, we compared the purification yield of VRC and Mt. Sinai spike from transiently transfected cells incubated at either 32°C or 37°C after the addition of enhancers 18-hours post-transfection. As seen in **Table 1** and **Fig. 2B**, the lower temperature expression led to a dramatic increase in yield to ∼5 mg/l for both CoV-2 spike proteins we tested. This yield is similar to that cited in a recent report of the Mt. Sinai spike protein produced at 37°C and using a diafiltration approach to buffer exchange [3]. However, unpublished results from other colleagues in the field using this construct suggest 1-2 mg/l is a more consistently observed result, suggesting that these improvements will make a marked enhancement to yield. Published VRC spike protein yields are 0.5 mg/l [2], suggesting that our modified procedures improved production of this protein by nearly 10-fold.

We also investigated the time of harvest as a variable, as this is a common approach to improve yield from secreted protein platforms [11]. In initial experiments incubated at 37°C, increasing the time of harvest from 72 hours post-transfection to 96 hours resulted in some improvement in yield (**Table 1**). We did not test production at the lower temperature of 32°C at 72 hours, as cell growth is considerably slower at this temperature and we anticipated this would not be enough time for high levels of expression. Therefore, 96 hours was used as the standard for 32°C expression as noted above. However, we did explore longer growth times at the lower temperature, and saw no further yield increase for either VRC or Mt. Sinai spike by harvesting 120 hr post-transfection (**Table 1, Fig. 2B**). For these reasons, we chose 32°C and 96 hours as our optimal conditions for future expression.

Of note, we have also expressed and purified the analogous recombinant spike proteins from 4 other beta-coronaviruses using the original protocols described here as controls for the serology assay [4]. Interestingly, our yields of the spike proteins from SARS-CoV, MERS-CoV, OC43, and HKU1 (4.6, 10.6, 8.4, and 5.7 mg/l, respectively) in a single experiment from cells incubated at 37°C were considerably higher than that of the CoV-2 spike under similar conditions. This argues that SARS CoV-2 spike is unique in some aspect of protein stability and/or recombinant protein expression, as these other coronavirus spike proteins were cloned in the same vector and using the same strategy as the VRC CoV-2 spike. We anticipate that the new protocols outlined here might further improve the yield of those proteins as well.

### 3.3 Mode of target capture

Our observations of protein loss during steps in which spike is either concentrated or exposed to large surface areas, suggested that we might further improve purification by eliminating the TFF, which is necessary to achieve maximum protein capture from the culture supernatant during subsequent column IMAC. Thus, we used magnetized IMAC beads to capture VRC spike in batch mode from filtered lysates. The final protein purified with this approach was similar in terms of quantity and final purity (**Table 1** and **Fig. 2C**) to proteins purified by the TFF/IMAC/desalting protocol. This batch process has several distinct advantages over the more complex protocol. First, it eliminates the need for the labor-intensive TFF process, which for large-scale transient cultures can take many hours, and the requirement for specialized equipment and consumables. Second, when combined with a gravity flow approach to buffer exchange, this process completely obviates the need for a protein workstation, making the process accessible to many laboratories without FPLC technology, and making it highly scalable.

### 3.4 Quality control validation of spike proteins

We evaluated the functionality of spike proteins by structural and conformation-based methods. To assess the oligomeric state, spike proteins were analyzed by electron microscopy and analytical SEC (AnSEC). Negative-stain TEM images of spike particles **(Fig. 3A)** clearly show the expected trimeric structure and closely resemble previous published images of trimer spike proteins [2]. In addition, as seen in **Fig. 3B**, recombinant spike proteins eluted during AnSEC at a volume expected for a protein of ∼520 kDa rather than that expected for monomeric spike (∼180 kDa). This result was consistent over all preparations of both VRC and Mt. Sinai spike proteins, and in no cases did we observe any detectable monomeric spike protein by AnSEC. These macromolecular based approaches support the conclusion that our purified spike proteins adopt tertiary and quaternary structures as expected of SARS CoV-2 spike.

**Figure 3.**
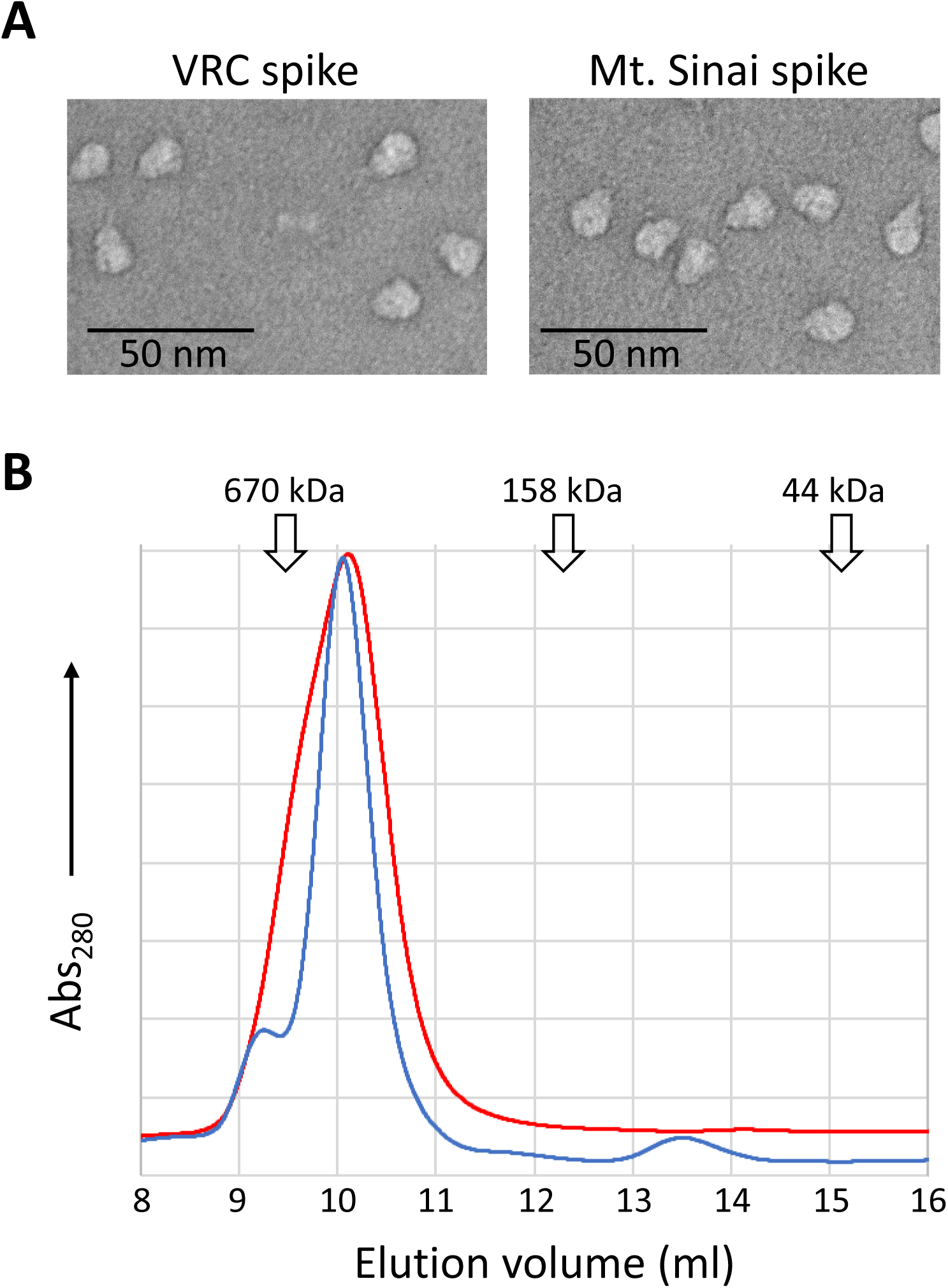
Assessment of the oligomeric state of VRC and Mt. Sinai spike proteins. **A.** Representative negative-stained transmission electron micrographs of VRC and Mt. Sinai spike proteins. **B.** Analytical size exclusion chromatography of purified VRC (blue line) and Mt. Sinai (red line) spike proteins. Peak elution volumes of sizing standards are noted (670 kDa - thyroglobulin, 158 kDa – γ-globulin, 44 kDa – ovalbumin).

An initial lot of VRC spike protein was previously used to develop and optimize a serology assay using an ELISA format [4]. To assess whether spike proteins produced by our modified protocols had equivalent ELISA sensitivity, multiple production lots of VRC spike were compared in the serology assay. **Fig. 4** demonstrates that independent batches of VRC spike proteins performed nearly identically in the assay at multiple concentrations of positive control sera. Assay sensitivity was consistent across multiple lots of protein produced with the same expression conditions (duplicate bars of identical color) and also across protein produced at different expression times and temperatures. Consistent lot-to-lot performance of spike protein is an essential part of a high-quality, sensitive serology assay.

**Figure 4.**
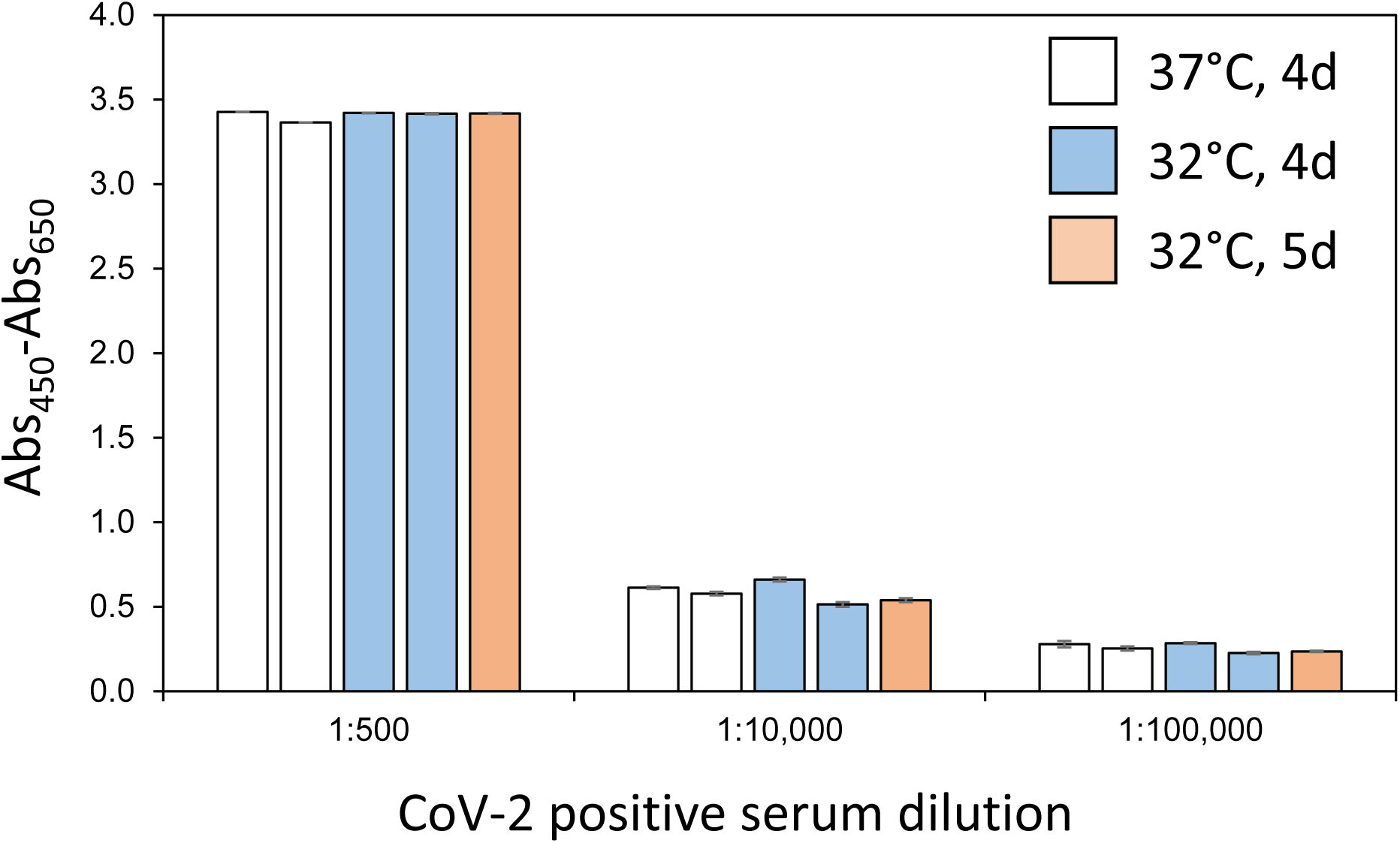
ELISA sensitivity of selected VRC Spike production lots. Multiple lots of VRC spike proteins generated using the noted conditions of expression time and temperature were used to coat ELISA plates which were then treated with positive control patient sera at the indicated dilutions. All measurements were performed in triplicate and means are plotted with standard deviations noted with error bars. Measurements are based on absorbance at 450 nm internally corrected by subtraction of absorbance at 650 nm.

## 4. Conclusions

In summary, we presented multiple improvements to the production of SARS CoV-2 spike protein that will allow labs with modest protein expression and purification experience to consistently produce high-yield and high-quality protein for use in a variety of applications. While we optimized the process to generate proteins for serology assays, the high quality of these proteins makes them amenable for biochemical, biophysical, and structural studies, or as substrates for drug screening. We hope that the parameters we highlighted in this report will help others to overcome bottlenecks in spike production and guide future optimization work.

## Abbreviations

AnSEC: analytical size exclusion chromatography,
CV: column volume,
ELISA: enzyme-linked immunosorbent assay,
IMAC: immobilized metal ion affinity chromatography,
MWCO: molecular weight cut-off,
RBD: receptor binding domain,
SDS-PAGE: sodium dodecyl sulfate-polyacrylamide gel electrophoresis,
TEM: transmission electron microscopy,
TFF: tangential flow filtration,
VRC: Vaccine Research Center

## Acknowledgements

We acknowledge Dr. Matthew Hall (NIH/NCATS) and Dr. Matthew Memoli (NIH/NIAID) for their role in the development of the serology assays used to validate these proteins. The authors would like to acknowledge Kizzmekia Corbett and Olubukola Abiona of the NIAID VRC for their generous donation of coronavirus spike plasmids and technical support during our initial purification of spike proteins. This research was supported in part by the Intramural Research Program of the National Institute for Biomedical Imaging and Bioengineering. This project has been funded in whole or in part with Federal funds from the National Cancer Institute, National Institutes of Health, under contract number HHSN261200800001E. The content of this publication does not necessarily reflect the views or policies of the Department of Health and Human Services, nor does mention of trade names, commercial products, or organizations imply endorsement by the U.S. Government.

